# Human sensorimotor resting state beta events and 1/f response show good test-retest reliability

**DOI:** 10.1101/2023.08.16.553499

**Authors:** Amande Pauls, Pietari Nurmi, Heidi Ala-Salomäki, Hanna Renvall, Jan Kujala, Mia Liljeström

**Affiliations:** BioMag Laboratory, HUS Medical Imaging Center, Helsinki University Hospital, 00290 Helsinki, Finland; Department of Neurology, Helsinki University Hospital and Department of Clinical Neurosciences (Neurology), University of Helsinki, 00029 Helsinki, Finland; Department of Neuroscience and Biomedical Engineering, School of Science, Aalto University, 02150 Espoo, Finland; Department of Psychology, University of Jyväskylä, 40014 Jyväskylä, Finland

## Abstract

Neurological conditions affecting the sensorimotor system have a profound impact on individuals’ physical independence and are associated with a considerable socioeconomic burden. Reliable functional biomarkers allowing early diagnosis of these conditions or targeting treatment and rehabilitation can reduce this burden. Magnetoencephalography (MEG) can non-invasively measure the brain’s salient rhythmic patterns such as the somatomotor (‘rolandic’) rhythm. This rhythm shows intermittent high amplitude ‘events’ in the beta (14-30 Hz) frequency range which predict behavior across tasks and species and are altered by neurological diseases affecting the sensorimotor system. Thus, the sensorimotor resting beta phenotype is a promising candidate biomarker of sensorimotor function. A prerequisite for use as a biomarker is that it can be quantified reliably across different measurement sessions. Here, using MEG, we assessed the test-retest stability of spontaneously occurring sensorimotor power spectral characteristics, including both aperiodic (1/f) as well as beta band fluctuations (‘beta events’) in a cohort of 50 healthy human controls. Test-retest reliability across two separate measurement sessions was assessed using the intraclass correlation coefficient (ICC). Beta events were determined using a thresholding-based approach on a narrow-band filtered amplitude envelope obtained using Morlet wavelet decomposition across a range of parameters (recording length, amplitude threshold and filtering bandwidth). We find that both aperiodic power spectral features as well as several beta event characteristics show good to excellent testretest stability. Especially aperiodic component power spectral features (ICC 0.77-0.88), but also measures of beta event amplitude (ICC 0.74-0.82) were found to be very stable, while measures of individual beta event duration were less reliable, especially for the left hemisphere (ICC right ∼0.7, left ∼0.55). Recordings of 2-3 minutes were sufficient to obtain stable results for most parameters. Important for potential clinical applications, automatization of beta event extraction was successful in 86 % of cases. Beta event rate and duration measures were more sensitive to analysis parameters than the measures of event amplitude. The results suggest the sensorimotor beta phenotype is a stable feature of an individual’s resting brain activity even for short, 2-3 minute recordings which can be easily measured in patient populations, facilitating its use as a potential clinical biomarker.

## Introduction

Some neurologic diseases affecting the motor system, such as e.g. Parkinson’s disease, are difficult to diagnose at their early stages due to lack of easily observable brain structural changes such as e.g. brain MRI changes. Furthermore, disease trajectories and rehabilitation outcomes are variable and often unpredictable. Currently, biomarkers for estimating individual disease courses are lacking. Functional biomarkers reflecting the processes underlying motor dysfunction might help in the differential diagnostics, or in estimating, e.g., the rate of disease development or the recovery potential in individual patients. Such markers could also improve targeting of treatment and rehabilitation.

Non-invasive electrophysiological recordings, such as electroencephalography (EEG) and magnetoencephalography (MEG), measure brain activity resulting from the spatial and temporal summation of cellular neural activity of the brain’s underlying cortical areas (Buzsáki et al., 2012). The measured activity depends on factors such as neuronal density, size and shape, the anatomy of neural network connections, and their relative activity at any given point (Buzsáki et al., 2012). Thus, MEG and EEG measures reflect the effect of different structural and functional changes in cortical activity and have considerable potential as functional biomarkers.

One promising candidate functional biomarker is the rolandic 20-Hz beta rhythm which is observed consistently in humans (Hari and Salmelin, 1997) and across other species (Feingold et al., 2015; Haegens et al., 2011; Sherman et al., 2016). Cortical beta activity plays an integral role in several perceptual and cognitive functions and it is modulated in a variety of tasks including tactile processing (Haegens et al., 2011; Pfurtscheller et al., 2001), movement (Feingold et al., 2015; Salmelin and Hari, 1994), action perception (Babiloni et al., 2002; Hari et al., 1998) and attention (Sacchet et al., 2015; Van Ede et al., 2011).

Cortical beta band activity displays a characteristic pattern of bursting over time, occurring in intermittent high amplitude ‘beta events’ alternating with lower amplitude periods (Feingold et al., 2015; Jones, 2016). Beta activity is particularly patterned in the sensorimotor cortex (Seedat et al., 2020), where beta event rate predicts behavioral outcome in humans and rodents across tasks (Shin et al., 2017). In humans, spontaneous resting EEG beta band power (Smit et al., 2005; Van Beijsterveldt et al., 1996), as well as beta event parameters (Pauls et al., 2023) have been shown to be heritable.

Neurological conditions with related motor dysfunction, such as stroke (Bartur et al., 2019; Laaksonen et al., 2013, 2012; Parkkonen et al., 2018; Rossiter et al., 2014; Schulz et al., 2021), Parkinson’s disease (Pauls et al., 2022; Vinding et al., 2020) and amyotrophic lateral sclerosis (ALS) (Dukic et al., 2022; Proudfoot et al., 2017) are associated with changes in the sensorimotor cortical beta band signal. Furthermore, sensorimotor beta characteristics correlate with symptom severity (Bartur et al., 2019; Laaksonen et al., 2012; Parkkonen et al., 2018; Pauls et al., 2022; Rossiter et al., 2014) and clinical recovery (Laaksonen et al., 2013, 2012; Parkkonen et al., 2018).

In addition to rhythmic, or ‘periodic’, components, spontaneous cortical activity also contains prominent aperiodic (‘1/f’) components which show exponential decay characteristics. This aperiodic signal is postulated to reflect excitation-inhibition balance (Gao et al., 2017), and it is modulated, e.g., by brain maturation (Hill et al., 2022; McSweeney et al., 2021; Tröndle et al., 2022) and aging (Voytek et al., 2015; Wilson et al., 2022). It is also highly heritable (Pauls et al., 2023) and appears to be altered in several neurological and neuropsychiatric conditions, such as Parkinson’s disease (Helson et al., 2023), dystonia (Semenova et al., 2021) and ADHD (Ostlund et al., 2021).

Taken together, these studies have shown that sensorimotor beta activity and aperiodic fluctuations are prominent, spontaneously occurring and heritable characteristics of ongoing brain activity that are closely linked to sensorimotor functions, are preserved across mammalian species and show changes associated with sensorimotor symptoms in different neurological disease conditions. Given their salience and ease of acquisition, they are thus potential cortical biomarkers of sensorimotor disease state and its reactivity to treatment.

However, a prerequisite for using them as biomarkers is their intraindividual signal stability. In general, test–retest reliability for various MEG responses of clinical interest is good, as suggested, e.g., for measures of somatosensory processing (Illman et al., 2022; Piitulainen et al., 2018), picture naming (Ala-Salomäki et al., 2021), as well as whole-brain spontaneous oscillatory power (Martín-Buro et al., 2016) and resting state functional connectivity (Garcés et al., 2016). Earlier EEG studies have also demonstrated good testretest reliability of global beta band power at rest (Fingelkurts et al., 2006; Pollock et al., 1991). Decomposing cortical sensorimotor activity into its different dynamic components, or ‘beta events’, adds detail compared to the assessment of mere beta power globally, and the different beta event parameters’ test-retest reliability has not been assessed before. Therefore, we determined whether and to what extent different spontaneous sensorimotor beta event parameters and aperiodic activity are reliable across sessions.

## Materials and methods

### Subjects

50 healthy subjects (age mean +/-STD 45 +/-20 years, range 21-70 years) screened to exclude pre-existing neurological disorders, learning disabilities, and language disorders were included in the study after giving written informed consent. The study was approved by the Aalto University ethics committee and carried out in accordance with ethical guidelines set out in the Declaration of Helsinki.

### MEG recordings

Measurements were performed in a magnetically shielded room (Imedco AG, Hägendorf, Switzerland) with a 306-channel Vectorview neuromagnetometer (Megin Oy, Helsinki, Finland) consisting of 204 planar gradiometers and 102 magnetometers. Spontaneous cortical activity was recorded with a 1 kHz sampling rate, continuous head position monitoring (cHPI) and band-pass filtering at 0.03-330 Hz in two separate sessions one-two weeks apart, for five minutes each, while participants were resting with their eyes open. Vigilance was assessed via video during measurements to control subjects kept their eyes open.

### MEG signal processing and parameter extraction

Subjects’ data were assessed visually for vigilance effects and to exclude artifacts and periods with significant artifact were excluded from further analysis. For suppressing external artifacts, MEG data were preprocessed using the temporally extended signal space separation method (tSSS, (Taulu and Kajola, 2005) implemented in the MaxFilter software (Megin Oy, Helsinki, Finland, version 2.2.15)). Subject’s head movements were compensated based on the cHPI recordings, and individual MEG recordings were transferred to a common head space using MaxFilter’s signal space separation-based head transformation algorithm. One subject was excluded because head transformation to the common space introduced considerable noise, compromising data quality. Further signal processing was done using MNE-python version 1.3 (Gramfort et al., 2013). After band-pass filtering the data to 2-48 Hz with a one-pass, zero-phase, non-causal FIR filter (MNE firwin filter using a Hamming window), power spectral densities (PSD) were calculated using Welch’s method with a non-overlapping Hamming window and 2048-point Fast Fourier transformation.

### Channel selection

The subsequent analysis steps are illustrated in **Figure 1** extending the approach previously used (Pauls et al., 2023, 2022). For each hemisphere, we defined a region of interest (ROI) of 15 gradiometer channel pairs per hemisphere centered over the sensorimotor cortices. To quantify the PSD at each recording site, we computed the vector sum of the two orthogonally oriented planar gradiometers at each sensor location (‘vector PSD’):

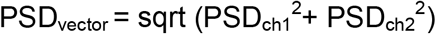

**Figure 1:**
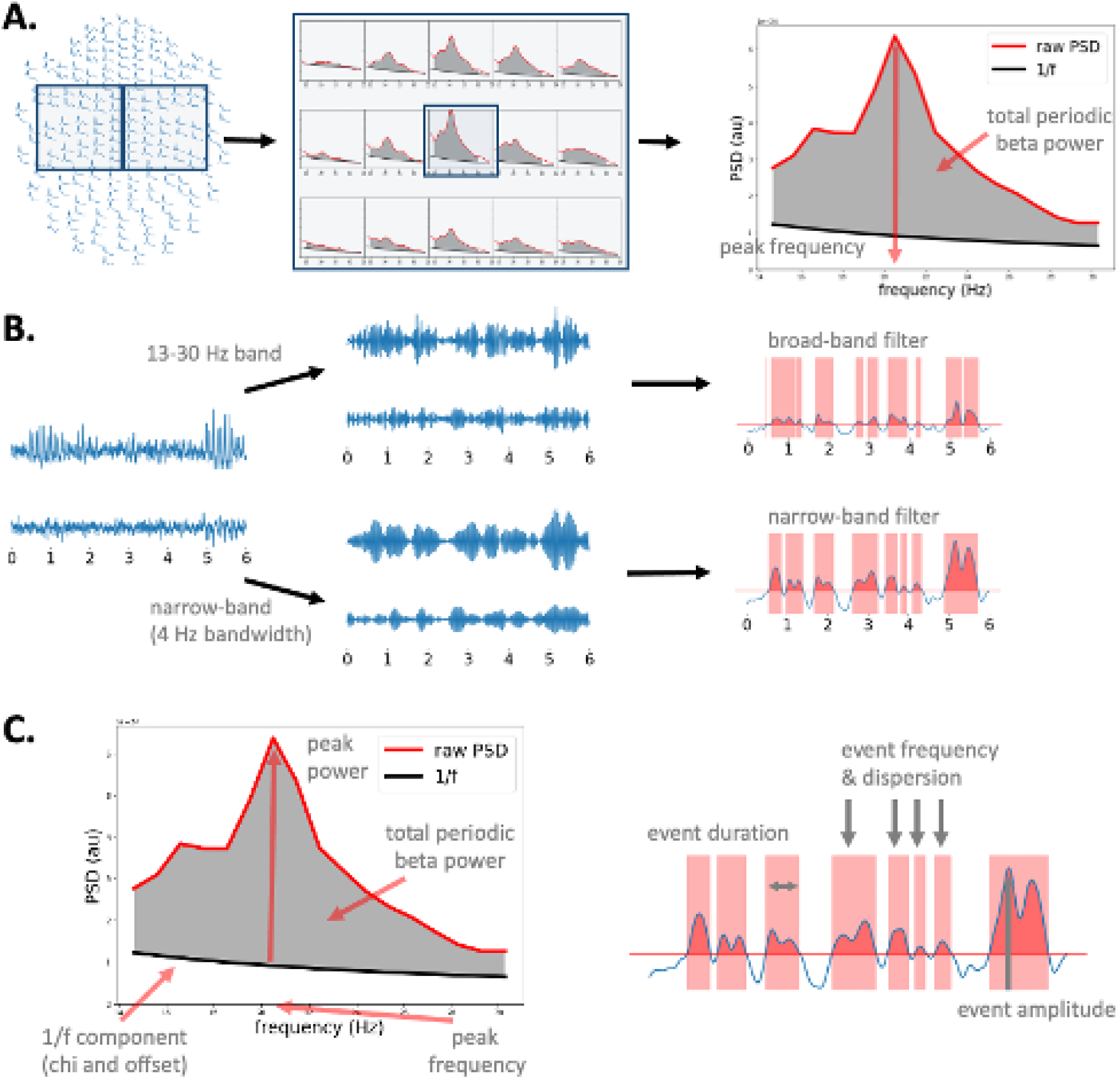
Sensorimotor phenotyping. A. Channel and peak frequency selection: In a predefined region of interest (ROI), channel pairs were combined using vector sum calculation, and the aperiodic (1/f) component was extracted using FOOOF as described previously (Pauls et al. 2022, Pauls et al. 2023). Peak channel and frequency within the ROI were selected either (1) manually by selecting the channel with the highest beta spectral peak frequency, (2) in an automated fashion by selecting the channel with the highest periodic beta power, or (3) using automation with visual-manual correction of the selected frequency and channel if necessary. **B**. Beta event extraction: The raw signal was convolved with a set of complex Morlet wavelets, and the resulting signal averaged either broad-band (14-30 Hz) or narrow-band (+/-2 Hz around the peak beta frequency) to obtain an amplitude envelope which was thresholded at the 75^th^ percentile (red line). Periods exceeding this threshold are defined as ‘beta events’. **C**. Sensorimotor phenotype parameters included PSD peak power and frequency in the 14-30 Hz beta band, total periodic beta power, 1/f exponent (chi) and offset, as well as beta event characteristics including event duration and amplitude mean, median, standard deviation, robust maximum, event rate and dispersion.

The resulting 15 vector-sum PSDs per hemisphere were then decomposed into a periodic and an aperiodic component using the FOOOF algorithm (Donoghue et al., 2020). After subtraction of the aperiodic component, the remaining periodic component was plotted for all 15 vector-sum PSDs. From the vector-sum PSD spectra, the channel pair with the most prominent spectral peak in the beta range was selected per hemisphere (‘the peak channel pair’) and the frequency of the power peak was noted (‘peak beta frequency’) (see **Figure 1A**). This choice of channel and frequency was carried out in three different ways:(1) an entirely automated approach, where the ‘peak channel’ was selected based on the area under curve (AUC) of the periodic part of the PSD between 14 and 30 Hz, and the peak was detected automatically as the highest amplitude in this frequency range; (2) a manual peak detection approach, where vector-sum PSD plots were visually inspected, the frequency of the highest peak noted, and the channel with the maximum amplitude at this frequency selected as the ‘peak channel’; and (3) a combined approach where peak channel and peak frequency were selected automatically as described, but all plots underwent visual control and the assignment of peak frequency (and sometimes channel) was re-adjusted if necessary. Typical reasons for correction of the peak frequency (and channel) were cases with strong alpha peaks and weak beta peaks, where the automatically detected beta peak was located on the shoulder of the alpha frequency peak.

### Beta event extraction

The channel pair and peak beta frequency corresponding to the chosen peak vector-sum PSD were used for beta event analysis (see **Figure 1B**). The channel pair’s raw, unfiltered time series data were downsampled to 200 Hz, high pass filtered at 2 Hz and decomposed by convolving the signal with a set of complex Morlet wavelets within the frequency range of 7-47 Hz with 1 Hz resolution and n_cycles=frequency/2. After this, the amplitude envelope was derived by averaging the signal within a certain beta frequency range. This was done either (1) broad-band across the entire beta frequency band (14-30 Hz) or (2) narrow-band, *i*.*e*., ± 2 Hz around the individual peak beta frequency chosen in one of the three ways described above. The vector sum over both channels’ beta band time series was calculated and rectified, to obtain one beta band amplitude envelope per channel pair. The envelope was smoothed with a 100-ms FWHM kernel and thresholded at the 75th percentile value. Periods exceeding this threshold for 50 ms or longer were defined as beta events. For event amplitude and event duration, the mean, median, robust maximum (defined as mean of the top 5% values) and standard deviation values were calculated (see **Figure 1C** for illustration). Furthermore, events per second (event rate) and event dispersion were calculated as described previously (Pauls et al., 2022). Times between beta events were defined as waiting times. To estimate the variation of waiting times (‘event dispersion’), we calculated the coefficient *C*_*V*_ proposed by (Shinomoto et al., 2005), defined as the waiting times’ standard deviation *σ* divided by their mean *μ*:

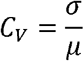

All values were calculated for both hemispheres in all subjects to obtain a sensorimotor signature phenotype (**Figure 1C)**. The sensorimotor phenotype features included PSD peak power and frequency in the 14-30 Hz beta band, total periodic beta power, 1/f exponent (chi) and offset, as well as beta event characteristics including event duration and amplitude mean, median, standard deviation, robust maximum, event rate and dispersion (see Figure 1C).

Beta event extraction was tested for a range of parameters to investigate their effect on the phenotype results and their stability. The narrow band filter bandwidth was varied (+/-1, 2, 3, 4, 5 Hz, and broad-band), and the amplitude threshold was tested for different percentiles (50th, 60th, 70th, 75th, 80th, 85th and 90th).

### Test-retest reliability analysis

Test-retest reliability was assessed using the intraclass correlation coefficient implemented in Pingouin (intraclass_corr function). ICC analyses were conducted on all the phenotype parameters described in **Figure 1C**, by comparing the outcomes between sessions 1 and 2. ICC is defined as follows:

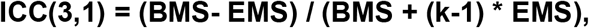

where BMS = between-subjects mean square, EMS = error mean square, and *k* = number of sessions. ICC below 0.4 is considered poor, 0.40-0.59 fair, 0.60-0.74 good, and 0.75-1.00 as excellent consistency (Cicchetti, 1994). We used ICC(3,1), with a fixed set of sessions (n=2) during each of which all phenotype parameters were assessed.

### Code and data availability

Data cannot be made publicly available due to Finnish data protection law. Data can, however, be shared for research collaboration with an amendment to the research ethics permit and a related data transfer agreement. All analysis code will be made available on GitHub.

## Results

### Test-retest reliability

Test-retest reliability was good or excellent for many of the phenotypic traits. **Figure 2** shows example scatter plots for three of the beta event parameters and their test-retest reliability in the left hemisphere.

**Figure 2.**
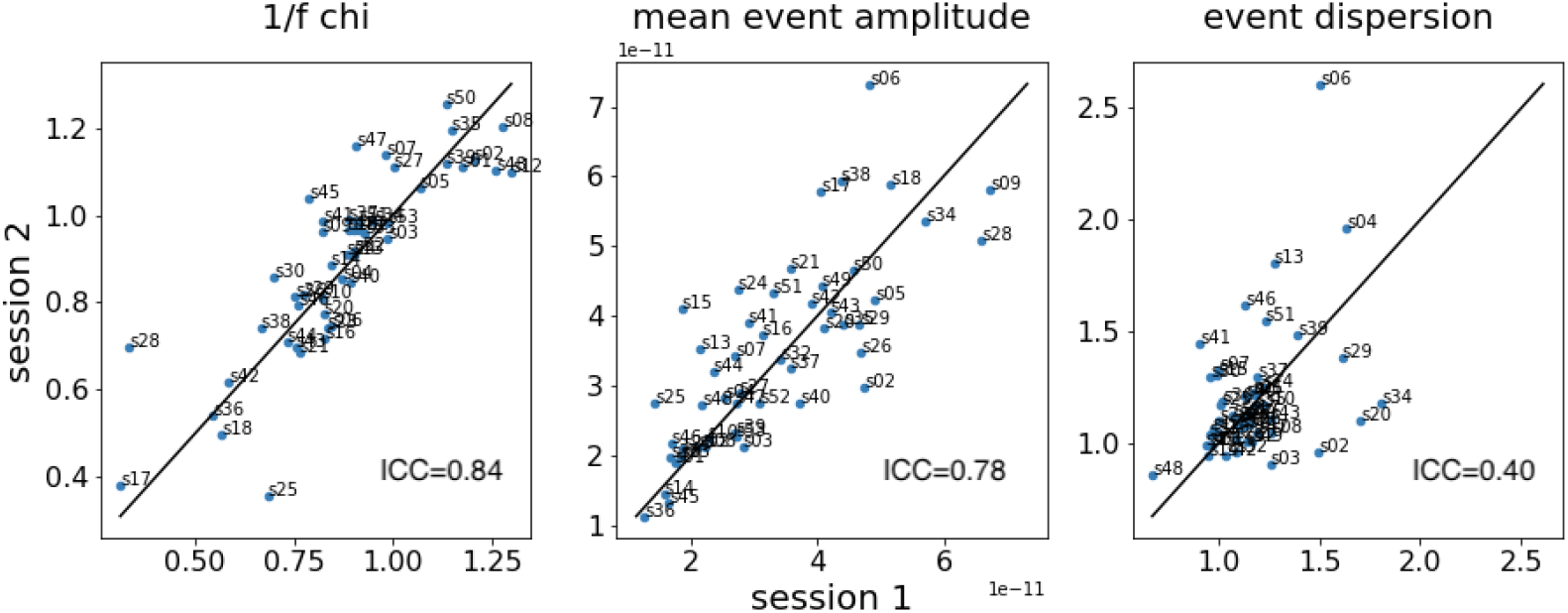
Scatter plots illustrating three of the sensorimotor phenotypes in the left hemisphere when using automated peak assignment with manual control. The points refer to individual subjects and subjects are labeled.

The three approaches using individual narrow-band filtering for determining beta events produced very similar results, whereas using a broad-band filter systematically altered the results (see **Figure 3**, discussed in more detail later). Entirely manual peak assignment did not significantly improve results and was very labor-intensive. On the other hand, automated peak assignment missed the PSD beta peak that was by visual evaluation the best one in 14 % of the cases. In most of these cases (86%), automatic peak assignment missed the periodic peak, usually in favor of a lower frequency peak located on the ‘shoulder’ of a large alpha frequency peak. In the remaining 14%, the peak channel changed because the periodic beta peak was visually more distinct in a different channel. We thus chose to work with the approach combining automated peak selection with manual control.

**Figure 3:**
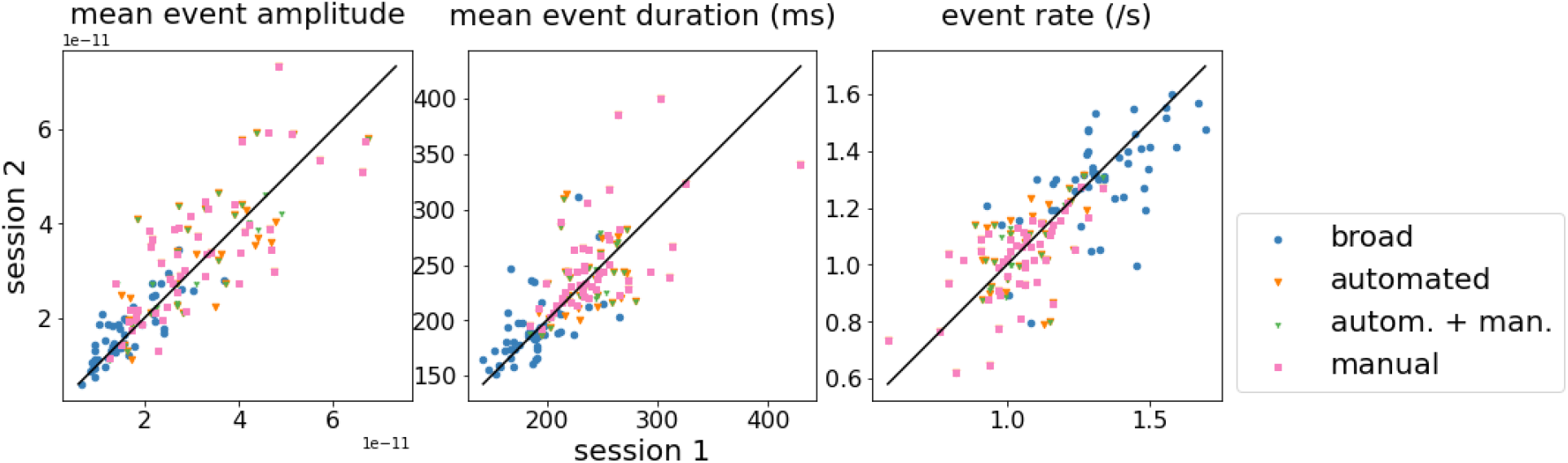
Effect of channel selection and event analysis strategies illustrated for different parameters (mean event amplitude, mean event duration and event rate). **Blue** – automated channel selection, broad band event extraction, **orange** – automated channel selection, narrow band event extraction, **green** – automated channel & peak selection and manual (human observer) control, **pink** – manual (human observer) channel & peak selection. Broad-band beta event extraction systematically shortens the duration of events, reduces their amplitude and increases their rate. Channel selection strategy has fewer and less systematic effects on the parameters.

Intraclass correlation coefficient (ICC) values for this approach (‘automated + manual’) are given in **Table 1**, and for all four approaches in **Supplementary Table 1**. Test-retest reliability for most parameters was good or excellent, but some parameters, notably event dispersion, proved poor (**Figure 2**). Interestingly, parameters of event amplitude were more reliable in the left hemisphere, whereas event duration parameters were more stable in the right hemisphere.

**Table 1:** Test-retest reliability of different sensorimotor phenotypes between two measurement sessions (automated peak selection with manual control). Light gray shading indicates ICC >=0.6, darker gray ICC >= 0.75.^*^The degrees of freedom df1 and df2 are 48 for all parameters.

|  | Left |  |  |  | Right |  |  |  |
| --- | --- | --- | --- | --- | --- | --- | --- | --- |
| PSD parameters | ICC | CI95 | F* | p value | ICC | CI95 | F* | p value |
| peak beta frequency | 0.90 | [0.83 0.94] | 18.7 | 4.322E-19 | 0.80 | [0.67 0.88] | 8.9 | 1.796E-12 |
| peak beta amplitude | 0.61 | [0.39 0.76] | 4.1 | 1.612E-06 | 0.69 | [0.51 0.81] | 5.5 | 1.482E-08 |
| 1/f chi | 0.84 | [0.74 0.91] | 11.7 | 8.131E-15 | 0.79 | [0.65 0.87] | 8.4 | 5.761E-12 |
| 1/f offset | 0.88 | [0.8 0.93] | 16.3 | 8.741E-18 | 0.77 | [0.63 0.87] | 7.9 | 2.113E-11 |
| total periodic beta power | 0.82 | [0.69 0.89] | 9.8 | 2.843E-13 | 0.86 | [0.76 0.92] | 13.2 | 6.779E-16 |
| Beta event parameters | ICC | CI95 | F* | p value | ICC | CI95 | F* | p value |
| duration mean | 0.57 | [0.35 0.73] | 3.6 | 7.892E-06 | 0.76 | [0.61 0.86] | 7.2 | 1.028E-10 |
| median | 0.57 | [0.35 0.73] | 3.7 | 6.978E-06 | 0.30 | [0.02 0.53] | 1.8 | 1.877E-02 |
| standard deviation | 0.59 | [0.37 0.74] | 3.8 | 3.754E-06 | 0.76 | [0.6 0.85] | 7.2 | 1.087E-10 |
| maximum | 0.41 | [0.15 0.62] | 2.4 | 1.491E-03 | 0.65 | [0.45 0.79] | 4.7 | 1.682E-07 |
| robust maximum | 0.59 | [0.37 0.74] | 3.9 | 3.514E-06 | 0.77 | [0.63 0.87] | 7.9 | 2.092E-11 |
| amplitude mean | 0.78 | [0.64 0.87] | 8.1 | 1.192E-11 | 0.76 | [0.61 0.85] | 7.2 | 1.072E-10 |
| median | 0.78 | [0.64 0.87] | 8.0 | 1.475E-11 | 0.75 | [0.6 0.85] | 7.0 | 1.859E-10 |
| standard deviation | 0.79 | [0.66 0.88] | 8.5 | 4.514E-12 | 0.76 | [0.61 0.86] | 7.3 | 8.184E-11 |
| maximum | 0.82 | [0.71 0.9 ] | 10.4 | 9.913E-14 | 0.74 | [0.59 0.85] | 6.8 | 3.212E-10 |
| robust maximum | 0.81 | [0.68 0.89] | 9.4 | 7.442E-13 | 0.76 | [0.61 0.86] | 7.3 | 8.706E-11 |
| event rate | 0.62 | [0.41 0.77] | 4.3 | 7.923E-07 | 0.79 | [0.65 0.88] | 8.5 | 4.750E-12 |
| event dispersion | 0.40 | [0.13 0.61] | 2.3 | 2.240E-03 | 0.31 | [0.03 0.54] | 1.9 | 1.522E-02 |

### Broad-vs. narrow band event characteristics

Use of broad-band filtering to determine beta event characteristics systematically affected event parameters, shortening event duration, increasing event rate and reducing event amplitudes compared to narrow-band event extraction (see also **Figure 3**). Furthermore, broad-band filtering decreased the signal to noise ratio (SNR), as illustrated in **Figure 4**.

**Figure 4:**
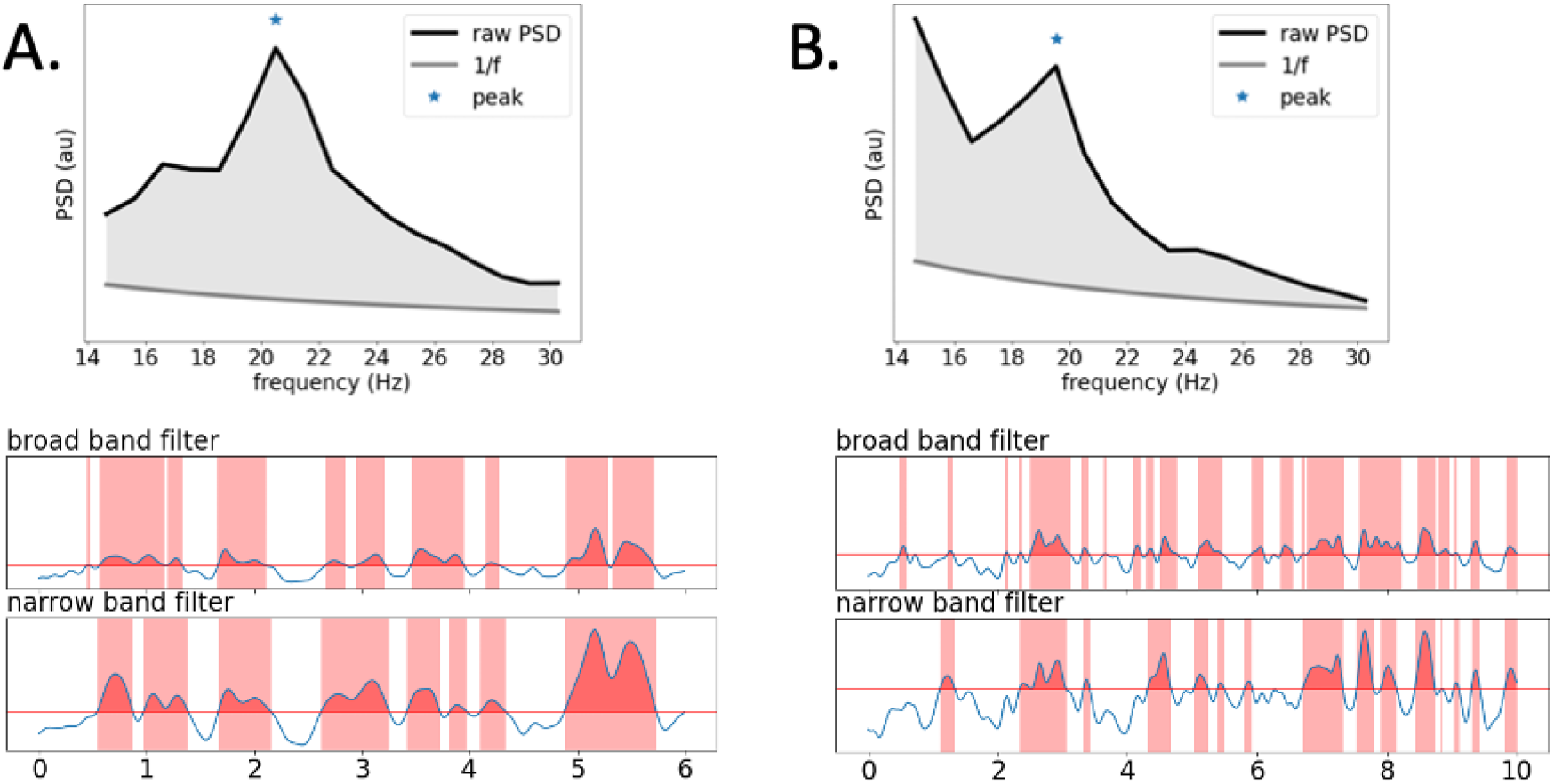
Effect of broad-band (14-30 Hz) vs. narrow band (peak frequency +/-2 Hz) filtering on amplitude envelope and signal to noise ratio (SNR), shown for a prominent (A) and a weaker (B) beta range spectral peak. The selection of the individual peak and bandwidth has relatively more effect in the prominent peak case in which the SNR decreases with increasing filtering bandwidth.

### Effect of recording length and processing parameters on ICC

Finally, we tested the effect of event extraction parameters (filtering bandwidth & amplitude threshold) as well as recording length on the test-retest reliability (see **Figure 5**). Overall, event amplitude showed good or excellent reliability and was relatively invariant to different parameters except for recording duration (**Figure 5**, middle column). Event duration was more noise- and parameter-sensitive (**Figure 5**, first column). Event rate reliability was fairly stable for longer recording durations. Event dispersion had the lowest reliability of all parameters, also at longer recording durations (**Figure 5**, last column).

**Figure 5:**
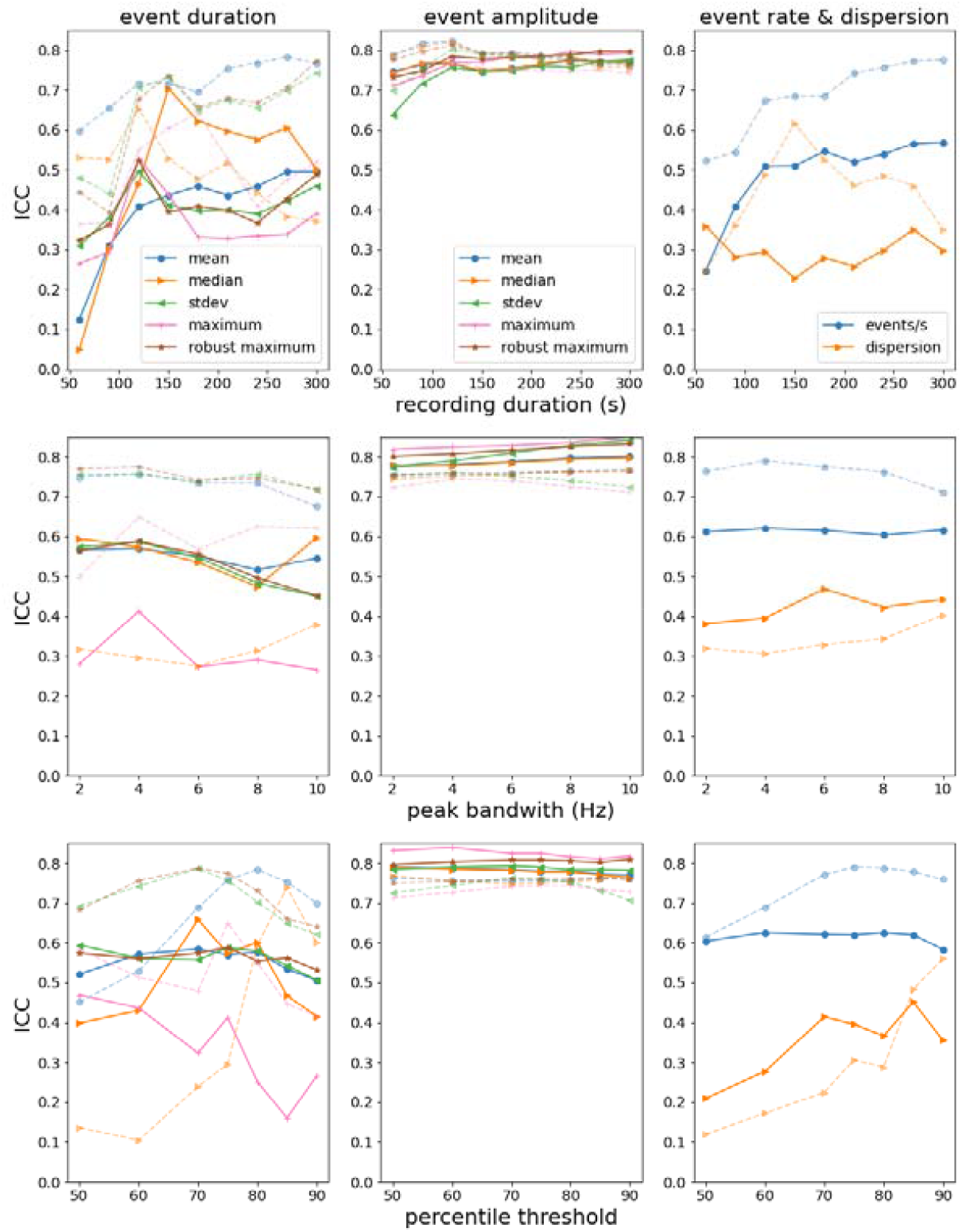
Effect of different event extraction parameters on session-to-session event parameter stability as assessed by ICC. Panels depict the following parameters: top row - effect of recording duration (s), middle row - peak bandwidth (Hz), bottom row - % amplitude threshold, with ICC on the y-axis. Columns: 1st - event duration, 2nd - event amplitude, 3rd - event rate & dispersion. Solid lines - left hemisphere, dashed lines - right hemisphere.

ICC stabilized reasonably well at 2-min recording length for most beta event parameters, so even short recordings may be enough for obtaining reliable results (**Figure 5**, top row). However, event rate estimates benefitted from longer recordings. A bandwidth of 4 Hz was optimal for event duration assessment as well as event rate, after which ICC decreased somewhat (**Figure 5**, middle row). Event amplitude appeared invariant to bandwidth. 70-80% percentile thresholds were optimal for both the event duration as well as the event rate parameter, while event amplitude was mostly invariant to this parameter (**Figure 5**, bottom row). Only dispersion appeared to benefit from higher percentile thresholds, raising ICC to around 0.5-0.6.

## Discussion

We demonstrate that human cortical sensorimotor dynamic cortical beta event parameters and 1/f characteristics as measured with resting-state MEG show good test-retest reliability. The results were robust across a range of analysis parameters including different filtering bandwidths and amplitude thresholds and appear to be stable even for just 2-3 minutes of recording for many parameters. The results suggest that sensorimotor beta phenotype is a stable feature of an individual’s brain activity with good potential as a clinical biomarker.

### Earlier studies of test-retest reliability of sensorimotor functional measures

Some previous studies have assessed sensorimotor system spontaneous and task-related measures’ test-retest reliability. In an MEG study assessing spontaneous resting-state oscillatory beta band power stability, ICCs ranged from 0.74 to 0.86 for frontal and parietal brain areas (Martín-Buro et al., 2016), comparable to our results for the left hemisphere beta power. In their study, beta power was separated into low and high beta power (13-20 vs. 20-30 Hz). Furthermore, the total band power included the aperiodic signal component, which was assessed separately here. Cortico-kinematic coherence, a measure of proprioception-related brain processing, has been demonstrated to have very good stability, 0.86 for the dominant and 0.97 for the non-dominant hand (Piitulainen et al., 2018). Stimulus-related sensorimotor beta suppression and rebound phenomena in response to sensory (tactile and proprioceptive) stimuli also show good to excellent stability (Illman et al., 2022). However, task-related functional measures rely on some degree of collaboration (and preserved function) from the subject, which can limit applicability in clinical settings. Thus, brief measurements of spontaneous brain activity as used here extend the spectrum of possible applications.

### Hemispheric differences

We found a degree of hemispheric lateralization for some of the parameters. The test-retest reliability was slightly higher for the left, dominant hemisphere for the amplitude parameter, probably reflecting the fact that left hemisphere spectra tend to have clearer periodic signal components. Interestingly, the beta event duration parameters as well as event rate were more reliable for the right hemisphere despite the fact that defining the peak was more difficult. A possible explanation for this result is that the dominant left hemisphere has more variable resting activity. On the other hand, the right hemisphere duration parameter may be more stable because of its relatively lower SNR, thus less reflecting the actual periodic beta signal fluctuations and more the general noise level or other, non-periodic activity. This effect can also be seen in the broad vs. narrow band signal extraction: broad-band signal extraction produces good ICC values, especially for the right hemisphere. The ICC difference between ‘broad’ and ‘narrow’ band extraction strategies is bigger for the left hemisphere with a more pronounced PSD beta peak.

### Potential of resting beta phenotype as a biomarker

There are no standard values determining acceptable reliability using ICC, but different suggestions have been made for their interpretation (Cicchetti, 1994; Koo and Li, 2016). ICC values above 0.6 (Cicchetti, 1994) or 0.75 (Koo and Li, 2016) have been considered to indicate good test-retest agreement, and values above 0.75 (Cicchetti, 1994) or 0.9 (Koo and Li, 2016) to indicate excellent test-retest agreement. Thus, many of the described event parameters in the present study show good to excellent test-retest reliability. A low ICC can relate to low test-retest agreement but can also relate to lack of variability among subjects (small dynamic range), small number of subjects or a low number of repetitions. The number of subjects in the current study should be sufficient to obtain reasonable ICC values and was the same for all studied parameters. However, the dynamic range was low, e.g., for event dispersion, with most subjects clustering in a very limited range of values, possibly contributing to the low ICC. Overall, the level of test-retest reliability obtained in the current study was good, supporting the use of the features of interest also in clinical settings.

Besides technical considerations, biological and pathophysiological considerations are also important for biomarker development. As outlined earlier, sensorimotor beta activity and dynamic beta events are detectable across different mammalian species including humans, non-human primates and rodents (Feingold et al., 2015; Haegens et al., 2011; Hari and Salmelin, 1997; Sherman et al., 2016) and they have been found to be heritable (Pauls et al., 2023; Smit et al., 2005; Van Beijsterveldt et al., 1996), suggesting that the brain’s sensorimotor signature is quite preserved across the sensorimotor system’s evolution. Furthermore, previous studies have shown disease-related beta changes at the group level (Bartur et al., 2019; Dukic et al., 2022; Laaksonen et al., 2013, 2012; Parkkonen et al., 2018; Pauls et al., 2022; Schulz et al., 2021; Vinding et al., 2020). Finally, the human brain’s sensorimotor system has been extensively studied and is quite well understood. We suggest that these factors, in combination with good test-retest reliability demonstrated here, make sensorimotor beta activity a good candidate electrophysiological biomarker.

### Limitations and future directions

Although the analysis approach used here was largely automated, it still required some manual preprocessing (adjustment of peaks in some cases with weak beta peak). The need for a human observer is always associated with a certain degree of observer bias, and different observers and experience levels can lead to increased levels of uncertainty. Furthermore, human observers need to be trained to make the process as reliable as possible. Here, only one observer (AP) carried out all manual beta peak selection. Future studies are needed to further automate beta event characterization.

Furthermore, sources of variations in signal to noise ratio (SNR) in PSD spectra need to be explored. Some subjects have relatively small periodic components in their PSD spectra for unknown reasons, and poor PSD SNR is not always clearly attributable to measurement noise. In many studies, subjects with poor SNR are excluded before further data analysis. However, after excluding clear problems with measurement quality (external noise), it would be interesting to explore reasons for poor SNR, which might in fact be related to brain processing features or brain state in these subjects. Here, we excluded one subject due to SNR considerations arising from problems related to data collection (poor head positioning during one session). Some subjects had bigger session to session fluctuations than others due to factors not obviously related to measurement factors. Future studies should assess the sources of session-to-session variability, e.g., factors such as vigilance. We assessed vigilance clinically during the measurement (via video) and the eyes open resting condition was used. The raw data was visually assessed for vigilance effects also (exclusion of gradual slowing of activity) to ensure steady vigilance levels. Thus, major fluctuations in vigilance have been excluded, but small changes in alertness or habituation effects across sessions are possible. Quantitative, automated vigilance assessment may also be helpful in the future.

The sensorimotor phenotype was assessed at the sensor-level. While source-level approaches add some level of spatial-anatomical resolution, they also introduce more data processing and analysis choices, making the approach less feasible for potential clinical applications and possibly leading to biased estimates. Furthermore, clinically useful source-level analyses would necessitate suitable cortical parcellations to avoid multiplying the amount of data. As MEG is most sensitive to sulcal brain activity, established parcellations taking this into account would be needed. Here, we were specifically interested in testing the reliability of a simple approach using sensor-level data to characterize the resting sensorimotor phenotype for potential clinical applications. If the potential caveats of source-level analyses (amount of data processing, automated analyses, suitable parcellations) can be solved, future studies should address whether more reliable measures can be obtained at the source-compared to the sensor-level.

### Pipeline recommendation

In the current study, recording durations of 2-3 minutes were sufficient to get stable beta event results for most parameters. For most of the examined parameters, short good-quality data segments were preferable to longer data segments with more variable data quality. Only the event rate, and to some extent the event duration parameters, benefited from longer recording times.

Automation of beta peak detection was successful for 86 % of the hemispheres: We recommend at least visual control to ascertain correct beta peak assignment. In the future, optimized automatization approaches which work for most subjects would be helpful to eliminate human observer bias. Alternatively, broad-band (13-30 Hz) beta event extraction can be done, but use of a specific beta peak channel and beta band peak for extraction of beta event information increased stability of the beta event duration parameter. A beta amplitude threshold of 75% and 4 Hz bandwidth appears appropriate, giving a good reliability for all beta event parameters. Estimation of event dispersion had low reliability but improved at higher percentage thresholds, so this parameter might require different processing settings.

### Conclusions

In summary, we demonstrate that a robust resting-state sensorimotor phenotype wth good or excellent test-retest stability can be obtained from MEG data in healthy subjects relatively easily even from short, 2-3 minutes long MEG recordings. This sensorimotor beta phenotype appears to be a relatively stable feature of an individual’s resting brain activity which can be easily measured also in patient populations, facilitating its use as a potential clinical biomarker.

## Acknowledgements & funding source

We thank all subjects for participating in the study. We acknowledge the following funding sources: AP received funding from the Academy of Finland (grant number 350242) and the Sigrid Juselius Foundation. Pietari Nurmi received funding from the Finnish Cultural Foundation and the Swedish Cultural Foundation in Finland. Heidi Ala-Salomäki received funding from the Jenny and Antti Wihuri foundation and the Finnish Cultural Foundation. HR received funding from the Academy of Finland (grant numbers 127401 and 321460). Mia Liljeström received funding from the Swedish Cultural Foundation in Finland.

