## Supplementary Table 1 for "Human sensorimotor resting state beta events and 1/f response show good test-retest reliability"

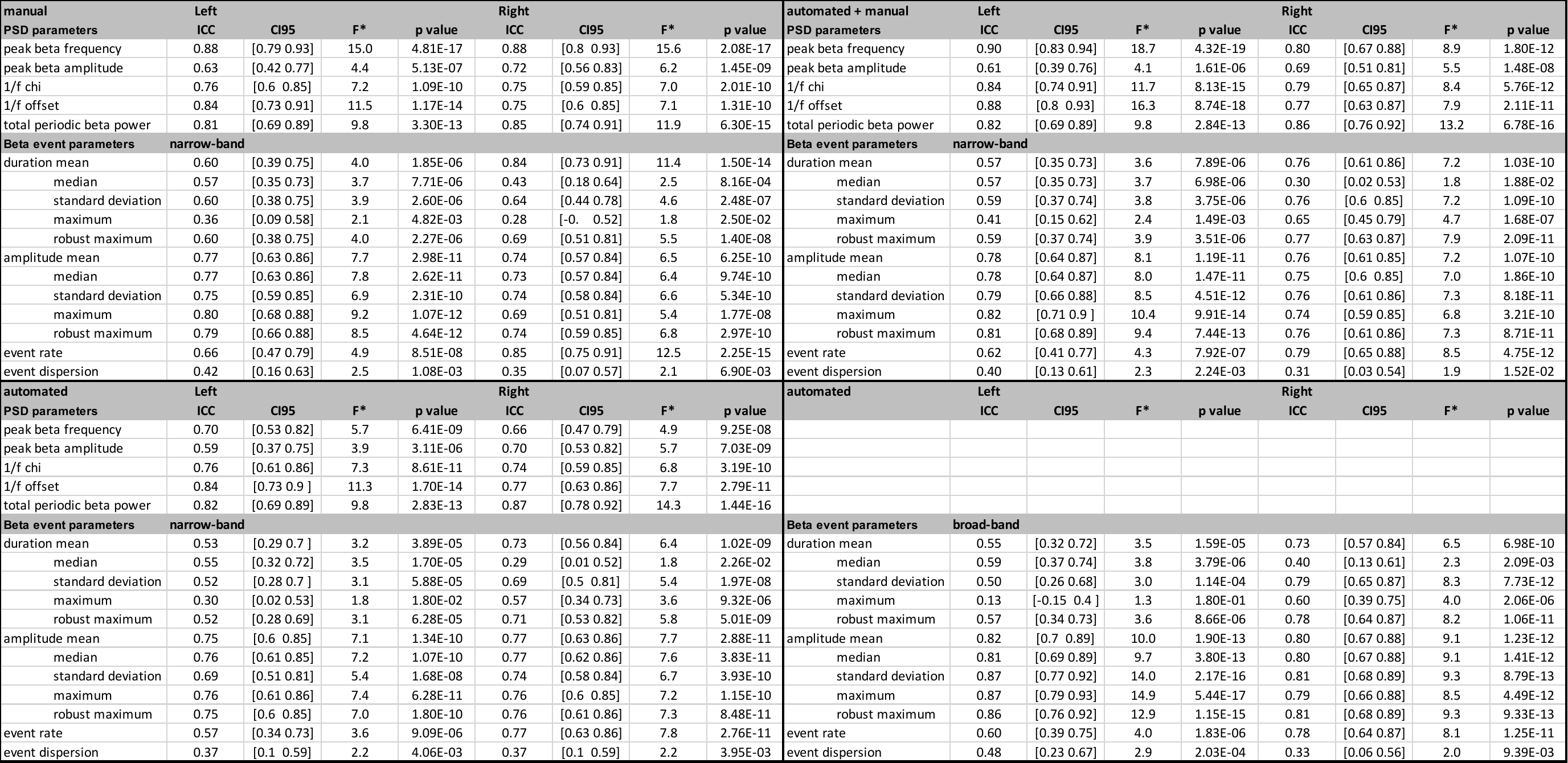

**Supplementary Table 1:** Test-retest reliability (assessed via ICC) of sensorimotor phenotype for all peak selection and event extraction approaches. *The degrees of freedom df1 and df2 are 48 for all parameters.
